# Eyeball translations affect saccadic eye movements beyond brainstem control

**DOI:** 10.1101/2022.08.24.505091

**Authors:** Johannes Kirchner, Tamara Watson, Jochen Bauer, Markus Lappe

**Affiliations:** University of Münster, Institute for Psychology, 48149 Münster, Germany; University of Münster, Otto-Creutzfeldt Center for Cognitive and Behavioural Neuroscience, 48149 Münster, Germany; Western Sydney University, School of Psychology, NSW 2751, Australia; University of Münster, Department of Clinical Radiology, 48149 Münster, Germany

## Abstract

Vision requires that we rotate our eyes frequently to look at informative structures in the scene. Eye movements are planned by the brain but their execution depends on the mechanical properties of the oculomotor plant, i.e, the arrangement of eyeball position, muscle insertions and pulley locations. Therefore, the biomechanics of rotations is sensitive to eyeball translation because it changes muscle levers. Eyeball translations are little researched as they are difficult to measure with conventional techniques. Here we study the effects of eyeball translation on the coordination of gaze rotation by high-speed MRI recordings of saccadic eye movements during blinks, which are known to produce strong translations. We found that saccades during blinks massively overshoot their targets, and that these overshoots occur in a transient fashion such that the eye is back on target at the time the blink ends. These dynamic overshoots were tightly coupled to the eyeball translation, both in time and in size. Saccades made without blinks were also accompanied by small amounts of transient eyeball retraction, the size of which scaled with saccade amplitude. These findings demonstrate the complex interaction between rotation and translation movements of the eye. The mechanical consequences of eyeball translation on oculomotor control should be considered along with the neural implementation in the brain to understand the generation of eye movements and their disorders.

## Introduction

Saccades are the most frequent movements we make and orchestrated in detail by dedicated brainstem circuitry [1]. They are initialized by a pulse phase, a brief burst in neural innervation to the agonist rectus muscle to produce rotational eye acceleration. Then, neural activity is reduced to an appropriate level to keep the eye steady at the desired gaze direction. Anatomical studies showed that each eye muscle has a connective tissue pulley that acts as a muscle lever in close proximity to the eyeball [2, 3]. Small changes in eyeball position or pulley location, therefore, have consequences for the biomechanics of rotational eye movements by changing the muscle levers. For example, gaze-dependent pulley locations have been proposed as a mechanical implementation of Listing’s Law [4], which pre-scribes torsional rotation as a function of gaze direction [5]. The pulleys of the horizontal rectus muscles are located only around 8 mm posterior to eyeball center, implying that even 1-2 mm of eyeball translation might be sufficient to have large implications on the kinematics of horizontal eye movements.

A strong translation movement of the eye occurs in conjunction with blinks. During a blink, the extraocular muscles simultaneously co-contract so that the eyeball as a whole is being lifted and retracted back into its socket [6]. Interestingly, other subsystems of oculomotor control are still able to operate while the eye is retracted. It has been shown that the period of eye occlusion during the blink can be used to correct fixation errors [7], anticipate target motion [8], reset torsion [9] and elicit saccades [10, 11]. The interaction between the blink and the saccade systems has received particular interest, because blinks inhibit omnipause neurons in the pons which also play a crucial role in the saccade premotor circuit [12–14]. Saccades that are made together with blinks are closely time-locked to blink onset and also slower than those without blinks [10, 11, 15, 16]. These altered saccade kinematics might be due to an interaction of blink and saccade systems in the brainstem or they could result from mechanical interference of the eyeball translation and the change in muscle levers. Observations of dynamic saccade overshoots during blinks [10, 16] hint at a mechanical contribution to the kinematics of within-blink saccades. Eyeball translation during blinks might affect the lever arm of the rectus muscles in such a way that a saccade that is is planned properly by the brainstem is deflected while the eye is retracted, thus producing the transient overshoot.

Dynamic properties of the oculomotor plant, like eyeball translation, are difficult to investigate with conventional eye tracking techniques. Recent advances in high-speed magnetic resonance imaging (MRI) allow obtaining anatomical image sequences of the eyes with sufficient spatiotemporal resolution to resolve saccadic eye movements. We used MREyeTrack, an eye tracking method allowing measurement of the kinematics of eye movements in terms of both translation and rotation by segmenting sclera, lens and cornea in each image [17]. Axial single-slice MRI data was acquired at a temporal resolution of 55.6 ms of both eyes from eight participants, who were instructed to make saccades with and without blinks. We aimed to study the relationship between eyeball retraction and the saccadic gaze trajectory, with a particular focus on investigating whether dynamic overshoots of within-blink saccades could be caused by eyeball retraction.

## Results

In three runs of dynamic MR data acquisition we presented two visual targets along the horizontal meridian at a distance of 5°, 10° and 20° from each other. Participants were instructed to continuously look back and forth between the two visual targets (Movie 1 - https://doi.org/10.6084/m9.figshare.20588178.v1). Then, we collected another three runs with identical stimuli but this time instructed participants to make the gaze shift while blinking. In order to determine the precision with which the MREyeTrack algorithm (Fig. 1a) is able to estimate gaze in our experimental setup, we compared motion estimations between left and right eye because each is determined independently by MREyeTrack. Horizontal gaze position was highly correlated between left and right eye (Pearson’s r >= 0.95 and p < 0.001) for all participants (Fig. 1b). The residuals from linear regression analysis had a mean standard deviation of 1.3°.

**Figure 1:**
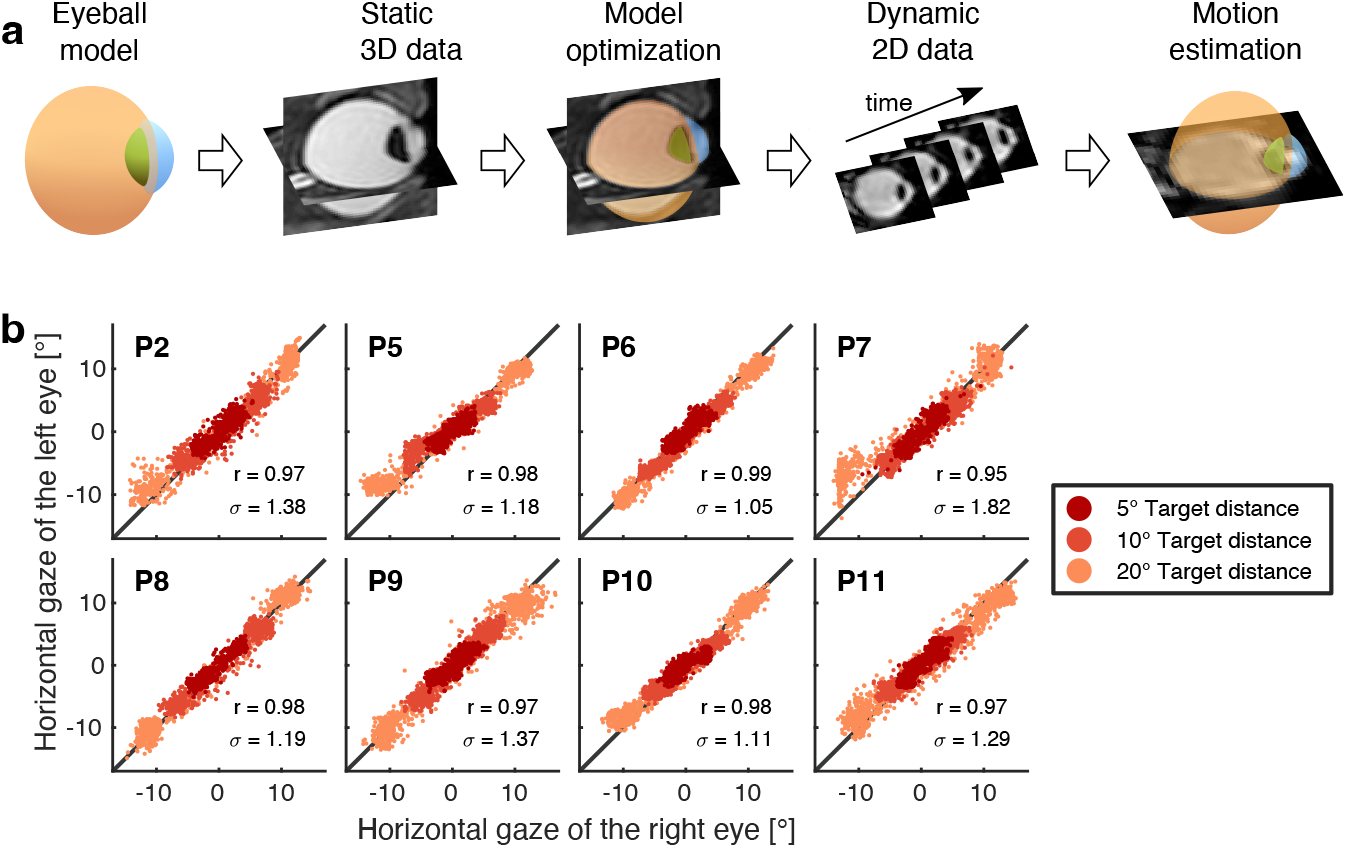
Eye motion estimation using MREyeTrack. **a** Illustration of the MREyeTrack workflow. The eyeball is modeled as a combination of three ellipsoids representing sclera, lens & cornea and subsequently optimized to the individual 3D MR data of each participant. Eye motion in the dynamic 2D MR data is then estimated by finding the optimal projection of the 3D eyeball model. Reprinted from Kirchner et. al [17]. **b** Precision of MREyeTrack gaze estimation by comparing left and right eye gaze while the participants looked back and forth (without blinking) between targets at three different distances. Pearson’s r of the linear regression (all p < 0.001) and standard deviation *σ* of the residuals are given for each participant.

### Dynamic overshoots of within-blink saccades

Saccades that are elicited within a blink sometimes show large dynamic overshoots in which the eye position far exceeds the target location but returns to the target location before the lid opens again (Fig. 2a & Movie 2 - https://doi.org/10.6084/m9.figshare.20588628.v1). Dynamic overshoots can reach several degrees of visual angle in size and occur right in the middle of a blink while the eye is retracted. In order to collect within-blink saccades of varying amplitude and direction, we instructed participants to shift their gaze, while blinking, between targets at 5°, 10° and 20° distance from each other. Occurrence of dynamic overshoots among the within-blink saccades was highly variable across saccade amplitudes and directions as well as across participants (Fig. 2b). While the within-blink saccades of one half of the participants (P5, P8, P9 and P10) showed barely any dynamic overshoots, the other half (P2, P6, P7 and P11) made them frequently. Their frequency of occurrence did not show any recognizable pattern with regard to saccade amplitude or direction. The size of dynamic overshoots also varied across participants and saccade properties (Fig. 2c).

**Figure 2:**
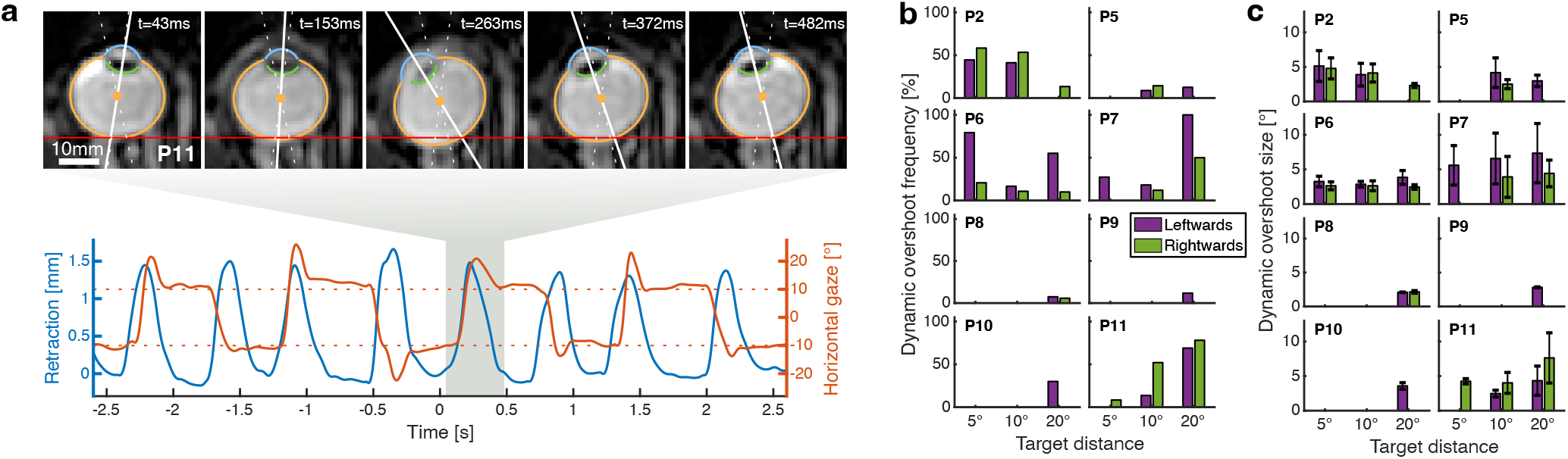
Dynamic overshoots of within-blink saccades. **a** Time series of horizontal gaze and retraction from 8 blinks, which can be identified by the gaussian-like peaks in the retraction data. Participants were instructed to shift their gaze between targets at +/- 10° (dotted horizontal lines) while blinking. These within-blink saccades often exhibited a dynamic overshoot. This is highlighted by the 5 MR images of one particular blink which show that gaze (solid white line) far exceeds the visual target (dotted white line) in the middle of the blink before returning near target position towards the end of the blink. Time is relative to onset of the highlighted blink. The location of the posterior eyeball border in the first image is marked in each image as a horizontal red line to illustrate the amount of retraction during a blink. **b** Frequency of dynamic overshoot occurrence from all within-blink saccades grouped into left- and rightwards saccades as well as target distance. **c** Same for the average size of the dynamic overshoots. Error bars are standard deviation.

### Binocularity of dynamic overshoots

The MRI recordings showed that dynamic overshoots were binocular with only little difference in size between the eyes (Fig. 3a & Movie 3 - https://doi.org/10.6084/m9.figshare.20589078.v1). In order to quantify overshoot binocularity, we compared the maximum gaze excursion between right and left eye for all within-blink saccades exhibiting a dynamic overshoot. These measures were highly correlated (Pearson’s r = 0.98, p < 0.001, Fig. 3b). The linear regression had an intercept of -1.8° degrees, suggesting an influence that is not related to the saccade. It might reflect the rotational trajectory of the blink-related eye movement, which is superimposed on all of the saccadic eye movement. The blink-related eye movement, albeit small, is of opposite polarity for the two eyes and might therefore be responsible for the intercept. We had measured blink-related eye movements for each eye in an earlier study with the same participants [6]. From this data, we calculated an averaged template from blinks without gaze shifts and subtracted this template from the within-blink saccade data of each eye. After subtraction, maximum gaze excursion of left and right eye were still highly correlated (Pearson’s r = 0.98, p < 0.001) and the intercept was reduced to -0.2° (Fig. 3c).

**Figure 3:**
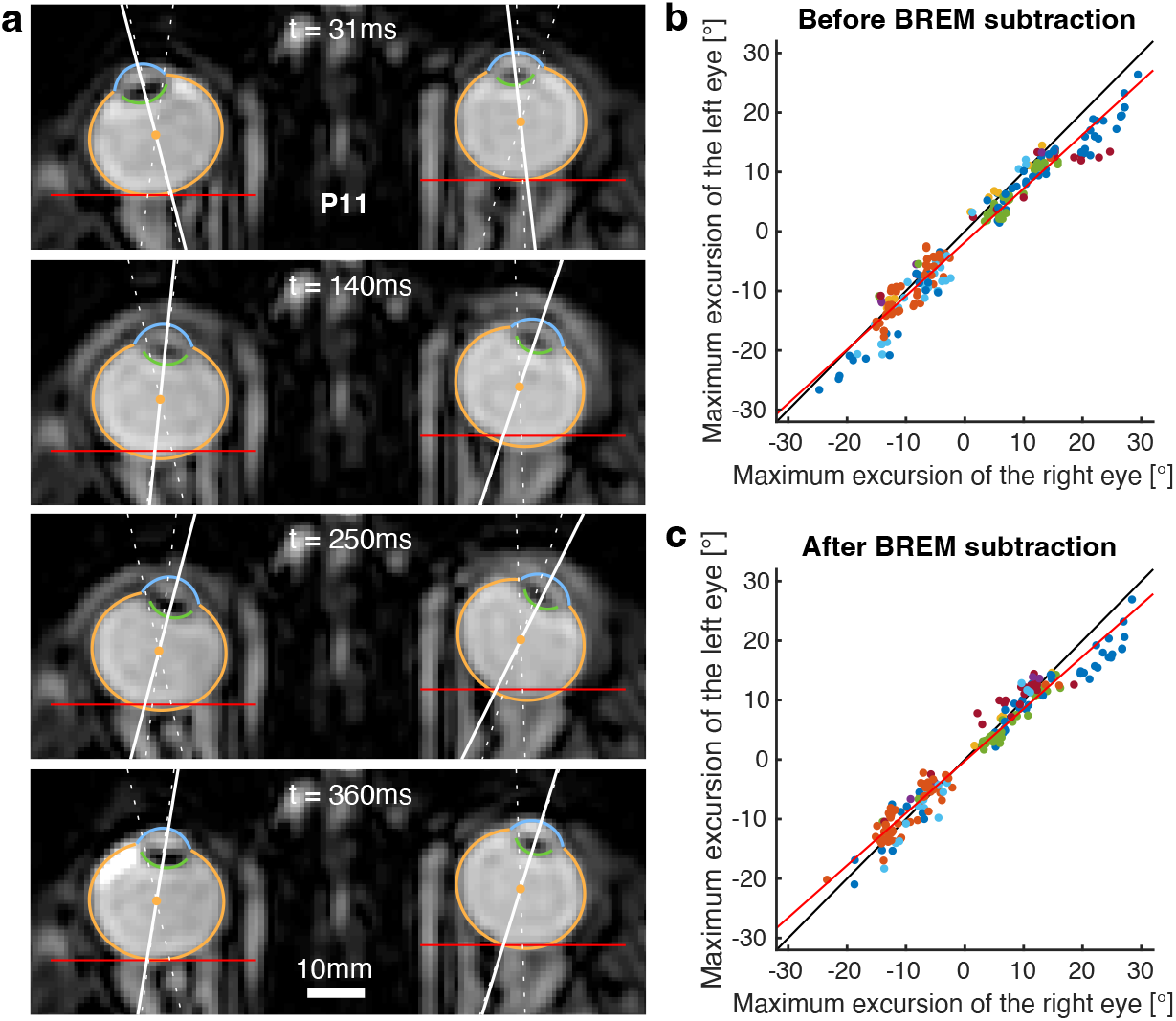
Binocularity of dynamic overshoots. **a** Example of a leftwards within-blink saccade with a dynamic overshoot. The third image from top shows the gaze position (solid white line) of both eyes moving beyond target position (dotted white line) before returning to the target in the fourth image. The location of the posterior eyeball border in the first image is marked in each image as a horizontal red line to illustrate the amount of retraction during a blink. Time is relative to blink onset. **b** Comparison of maximum gaze excursion during dynamic overshoots between left and right eye for all participants (each with a unique color) before subtracting the blink-related eye movement (BREM). Linear regression in red. **c** After subtraction of the BREM, the intercept of the linear regression is reduced to -0.2° from -1.8°.

### Relationship between eyeball retraction and dynamic overshoots

Dynamic overshoots may be caused mechanically by eyeball retraction or may be part of the neural signal that forms the saccade command. Crucially, if the saccade overshoot results mechanically from eyeball retraction, this would manifest itself in two ways. First, the overshoot would only occur after the retraction has reached full amplitude and is likely to be in close temporal proximity to the time of maximum retraction. Second, the return phase of the saccade overshoot would be an incidental consequence of the return motion of the retraction and should therefore be tightly time-locked to retraction return. In order to quantify these predicted relationships, we determined the temporal characteristics of the blink-induced eyeball retraction. We kept track of the time of maximum retraction and the time of retraction return, which we defined as the return movement reaching half-maximum retraction. Analogously, the temporal characteristics of the saccade trajectory where determined by overshoot and return, where saccade overshoot refers to the time of maximum gaze excursion and saccade return to the time when gaze is half-way between the overshoot and the final post-saccadic eye position (Fig. 4a).

**Figure 4:**
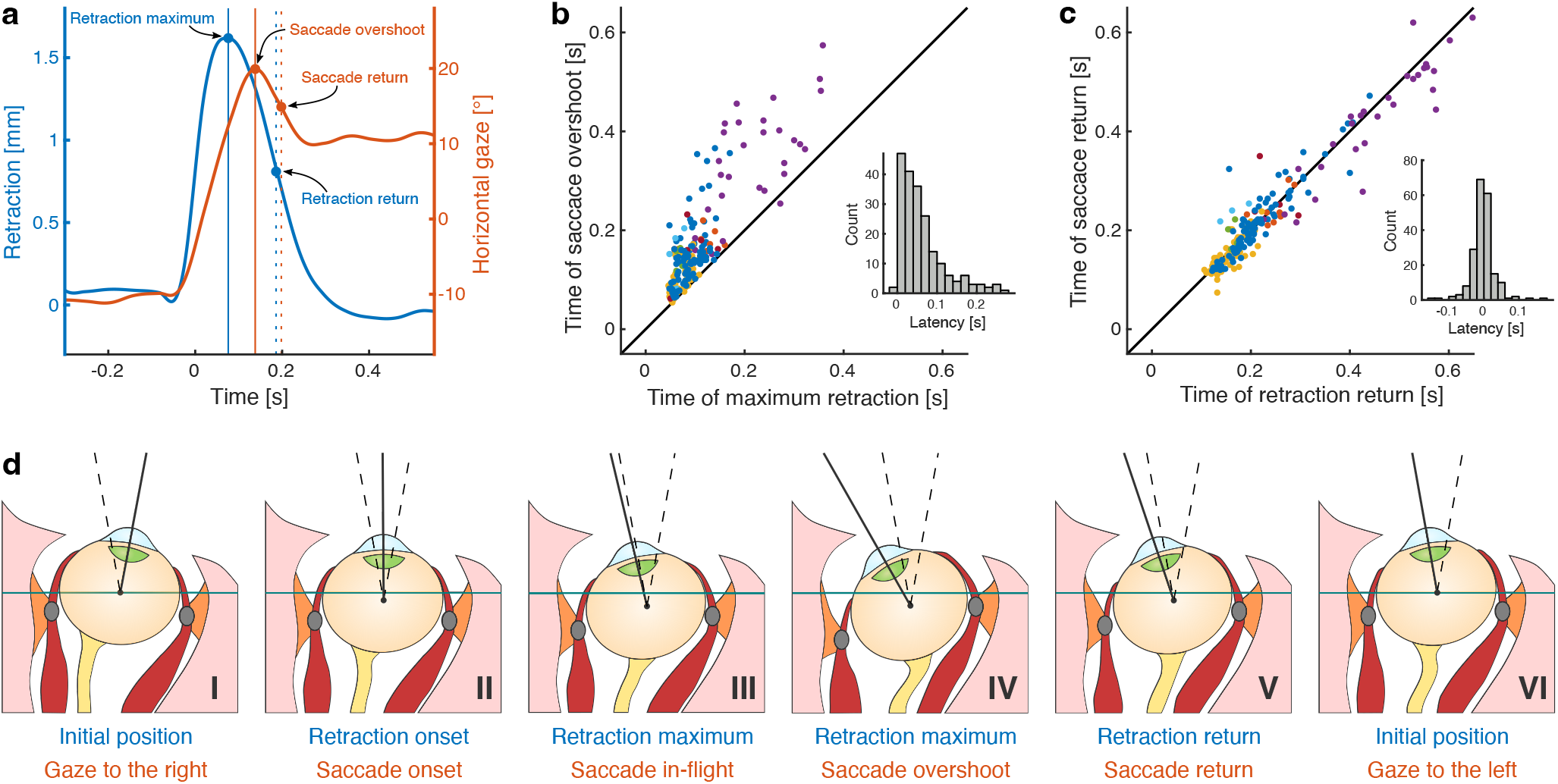
Relationship between dynamic overshoots and eyeball retraction. **a** Illustration of saccade and retraction metrics. The kinematics of eyeball retraction are defined by the time of maximum retraction, and the return, the time by which the retraction is at half-maximum. Analogously, saccade kinematics are defined by the saccade overshoot, the maximum gaze excursion, and the saccade return, which is the time by which gaze is half-way between overshoot and final position. **b** Time of saccade overshoot as a function of time of maximum retraction for all participants. Saccade overshoots occur always after retraction maximum and their latency follows a characteristic half-normal distribution. **c** Time of saccade return as a function of retraction return. The two measures are very tightly coupled to the point that a two-sided paired t-test shows no statistical difference. **d** Illustration of full eyeball kinematics during a within-blink saccades. Initially, gaze is directed on the right target (I). With blink onset, the eyeball retracts and the saccade is initiated shortly afterwards (II). While the eyeball reaches maximum retraction, the saccade is still in-flight (III), but reaches the overshoot peak soon after (IV). Return of the retraction and return of the saccade then follow the same time course (V), before the eyeball has returned to initial position and gaze is directed at the left target (VI).

We tested the relationship between these metrics for all within-blink saccades exhibiting a dynamic overshoot of at least 2° across all participants, which amounted to 204 saccades in total. For only 2 out of these 204 the saccade overshoot preceded the time of maximum retraction and in these two cases only by a few milliseconds (Fig. 4b). In all other cases, the saccade overshoot occurred after maximum retraction and followed a half-normal distribution with 50% of the overshoots occurring within 42 ms after retraction maximum. Next, we analyze d the temporal relationship between the time of retraction return and the time of saccade return, which we found to be very tightly correlated along the identity line (Fig. 4c). Subtracting the time of retraction return from the time of saccade return resulted in a normal distribution with a mean of 3 ms (SD = 34 ms), showing that the two measures follow an almost identical time course (Two-sided paired t-test, t(203) = -1.10, p = 0.27).

Based on these results, the sequence of events involving dynamic overshoots of within-blink saccades may be described as follows (Fig. 4d): Starting from initial gaze and eyeball position (I), the eyeball is being retracted into its socket with blink onset. This is then followed by the initiation of the saccadic eye movement shortly afterwards (II). The retraction reaches its maximum after around 100 ms, while the saccade is still in-flight (III). Global eyeball translation produces a change in orbital mechanics of the oculomotor plant such that the lever arm of the horizontal rectus muscles is changed. The neural innervation of the rectus muscles for a regularly-programmed saccade could then result in overextended gaze while the eyeball is still retracted (IV). The return motion of retraction and saccade then follow an identical time course (V), before the eyeball has returned to initial eyeball and final gaze position (VI).

### Factors contributing to dynamic overshoot size

Apart from the tight temporal coupling described above, we were interested to see to what extent eyeball retraction could also explain the size of dynamic overshoots. One factor should be the retraction amplitude, particularly if eyeball retraction leads to changes in orbital mechanics. To test whether this is the case we performed a median split analysis of the data in Fig. 5a. We grouped the saccade overshoot data in two groups having a retraction amplitude below or above the median of 1.19 mm. Indeed, the group with lower retraction amplitudes had a smaller overshoot size (1.65°, SD = 1.08°) than the group with higher retraction amplitudes (3.35°, SD = 3.11°, Two-sample t-test, t(496) = 8.16, p < 0.001). This implies that large retraction amplitudes are necessary for large dynamic overshoots to happen. However, large retraction amplitudes do not always lead to overshoots (Fig. 5a). This suggests that the occurrence of dynamic overshoots critically depends on the particular saccade kinematics and the relative timing to eyeball retraction. Even though the originally programmed saccade kinematics are not accessible from our data, it is still insightful to compare the timing of saccade overshoot relative to retraction return (Fig. 5b). Saccades that reached their overshoot before retraction return had an average overshoot size of 2.97° (SD = 2.68°), which was larger than that of saccades that reached their overshoot after retraction return (1.09°, SD = 0.61°, Two-sample t-test, t(496) = 7.74, p < 0.001). This demonstrates that the dynamic overshoots of within-blink saccades were larger if they occurred near to the maximum retraction.

**Figure 5:**
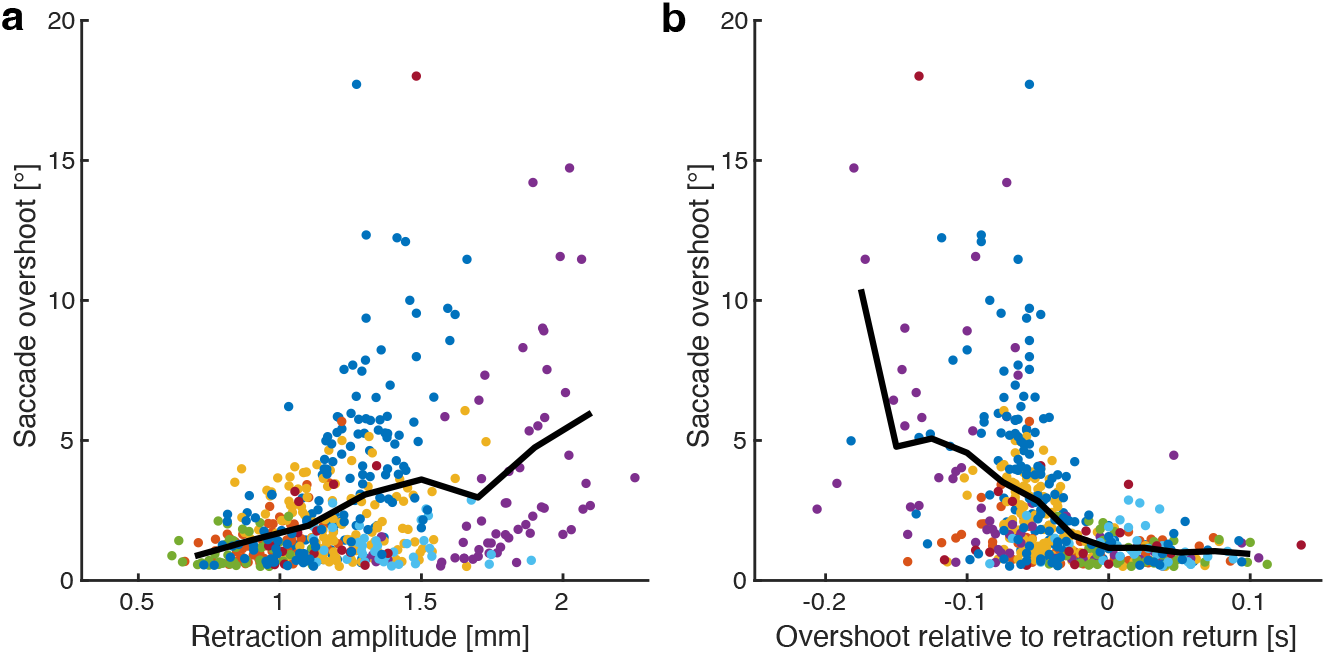
Factors contributing to dynamic overshoot size. **a** Saccade overshoot size as a function of retraction amplitude for all within-blink saccades with an overshoot of at least 0.5° from all participants. The black line is the moving average for bins of 0.2 mm width and showcases the positive correlation between retraction amplitude and saccade overshoot. **b** Saccade overshoot size as a function of the timing relative to retraction return. Overshoots which occurred several dozen milliseconds before retraction return and hence while the eye was fully retracted, were much larger than those occurring near or after retraction return. The black line is the moving average for bins of 25 ms width.

### Dynamic eyeball retraction of saccades without blinks

We finally also analyzed saccades without blinks. Even without blinks, the eyeball performs a small transient retraction following a horizontal saccade [18]. We wanted to test whether this retraction is directly related to the increased force exerted on the eye during the pulse phase of the saccade. If true, then the retraction should build up only while the saccade is in motion and the amount of retraction should scale with saccade amplitude. We investigated this in the saccades of 5°, 10° and 20° amplitude that were performed without blinks. There are two different sources of eyeball retraction present in this data. The first is a static retraction related to gaze direction [19]. The second is the dynamic retraction associated with the saccadic eye movement which we are interested in. It transiently decays after a few hundred milliseconds. To isolate the dynamic retraction, we averaged retraction trajectories across directions for all saccades of each target distance to remove the static translation. The resulting retraction trajectories clearly show the dynamic retraction associated with the saccadic eye movement (Fig. 6). Simultaneously with saccade onset, the retraction begins to build up and reaches its maximum just before saccade offset. After that, it slowly decays back to initial position. The pulse phase of the saccade coincides precisely with the time window during which retraction is building up and therefore suggests that the dynamic retraction is indeed caused by the increased force exerted by the rectus muscles. This is further corroborated by the fact that the amount of retraction scales with saccade amplitude. These dynamic retractions also occur during within-blink saccades, but we did not attempt to differentiate between blink- and saccade-related eyeball retraction because the blink-related retraction is a magnitude larger.

**Figure 6:**
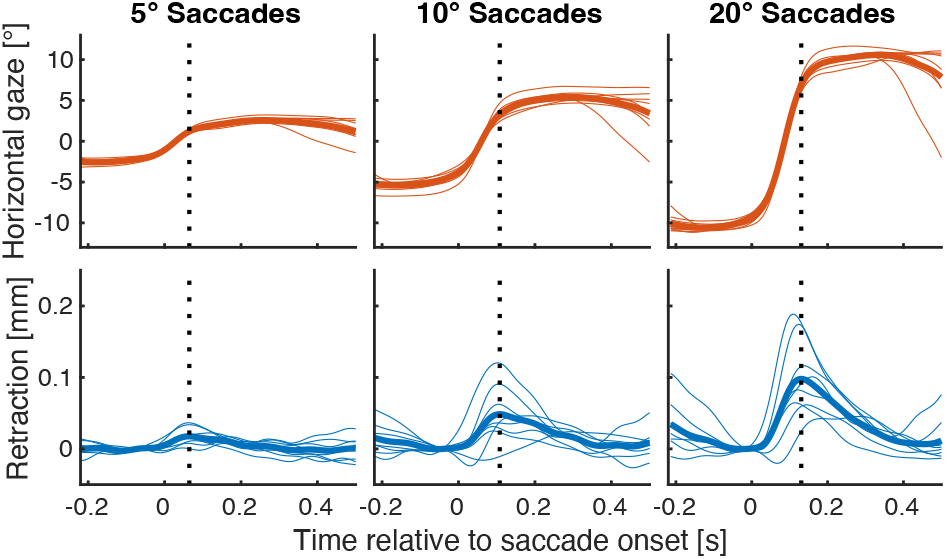
Dynamic eyeball retraction of saccades without blinks. Averaged gaze and retraction trajectories for each participant (thin solid lines) and across all participants (thick solid lines). The retraction reaches its amplitude (dotted vertical lines) once the acceleration phase of the saccade ends, after which it slowly returns to initial position. The amount of retraction scales with saccade amplitude and hence the force exerted by the horizontal rectus muscles on the eyeball.

## Discussion

We have shown that large transient overshoots occur when saccades are performed together with blinks, and that these overshoots can be attributed to the eyeball retraction that accompanies every blink. The timing of the saccade return motion after the overshoot was very tightly coupled to the return motion of the retraction, suggesting that the overshoot is not part of saccade programming in the brainstem, but instead an incidental consequence of eyeball translation. This is corroborated by our finding that the maximum amount of retraction explained much of the variety in overshoot size. Retraction alone, however, does not automatically cause the saccade to overshoot. Rather, retraction changes the properties of the oculomotor plant by affecting the pulleys and levers of the eye muscles such that a normal saccade command produces a different kinematic outcome. For vision, this is not a problem since the eyes are closed during the time of the overshoot. When the lid opens again the overshoot has already ceased. Saccade overshoots are likely the result of a coordination of eyeball retraction and a well-timed saccade command, such that the pulse phase of the saccade coincides with the retraction being near maximum. For saccades without blinks we also observed that the pulse phase led to small amounts of eyeball retraction which scaled with saccade amplitude. This demonstrates how closely rotational and translational motion components are connected in the execution of eye movements by the oculomotor plant.

How exactly could eyeball retraction change the properties of the oculomotor plant to generate dynamic overshoots? The torque needed to rotate the eyeball depends on the distance and angle at which the force exerted by the rectus muscles is applied, and therefore critically on the relative location of muscle pulleys to muscle insertion. At first glance, an eyeball retraction by 1 or 2 mm may not sound like it would change all that much, but horizontal rectus pulleys are located only around 8 mm posterior and the muscle insertion 8 mm anterior to eyeball center [2]. Therefore, eyeball retraction of 2 mm could lead to considerable change in the torque lever arm and hence the exerted force on the eyeball, even though the neural saccade command may not carry any overshoot signal. A similar scenario is encountered in the implementation of Listing’s Law, where the biomechanics of rotational eye movements are adjusted by gaze-dependent pulley locations. Because of this mechanical setup eye movements that are planned as 2D rotations in the in the superior colliculus [20] and the ocular motoneurons [21] can encompass the proper 3D torsional rotation implied by Listing’s law. Klier et al. demonstrated this directly by microsimulation of the abducens nerve, while the eye was initially in varying vertical positions [22]. Our data suggest that biomechanics are also responsible for saccade overshoot within blinks.

Inhibition of omnipause neurons in the brainstem can also lead to dynamic overshoots [1, 23], but is an unlikely explanation for the massive overshoots observed during within-blink saccades. Omnipause neurons gate the saccadic system by firing at a tonic rate, so that they need to be inhibited to trigger a saccade and resume firing once the saccade ends. Prolonged inhibition of omnipause neurons during blinks and a subsequently prolonged duration of the saccade pulse phase would produce overshoots. However, the saccades in our experiment reached their overshoot in the middle of the blink and returned to final gaze position while the blink was still on-going. Omnipause neurons are inhibited for the entire duration of the blink [24], so they cannot explain the occurrence of dynamic overshoots.

Our study provides a first step towards combining eye tracking with anatomical data of the oculomotor plant. The spatiotemporal resolution of our MRI data in combination with the MREyeTrack algorithm proved to be sufficient to resolve the trajectory of saccades between 5° and 20° amplitude and provided precise eyeball retraction data at the same time. However, the precision is not as good as that of conventional video-based eye tracking methods. For very detailed studies outside of blinks, for example for more detailed investigations into Listing’s Law, a hybrid approach of obtaining both eye tracking and MRI data might be fruitful. This would combine high precision data of the rotational eye motion trajectory with anatomical data of eyeball position or muscle pathway. For example, one could study muscle pathway deflections in the coronal plane with high-speed MRI while tracking rotational eye movements with a conventional eye tracker. This might be of particular interest for studying the mechanical implementation of Listing’s Law or eye movement disorder cases like strabismus. Although currently the direct assessment of pulley location is not possible with the data obtained in our study, more information of muscle path deflection due to eyeball lifting and retraction during blinks would further the understanding of dynamic overshoots in particular and of the oculomotor plant mechanics in general. A complete understanding of eye movements, from neural implementation in cortex and brainstem all the way to the periphery of the oculomotor plant, is essential to understand oculomotor control in health and disease.

## Methods

### Participants

Eleven healthy participants (P1-P11, age 23-49, 1 female, 10 males) participated in this study and gave informed consent. All procedures were approved by the ethics committee of the Department of Psychology and Sports Science of the University of Münster. The same participants took part in another study from our lab with identical experimental setup, but different experimental protocol and study goal [6].

### Experimental Setup

Eye movements were recorded in the axial plane at a temporal resolution of 55.6 ms using a 3T Philips Achieva Scanner (Philips Medical Systems, Best, The Netherlands) and a balanced steady-state free precession (bSSFP) MRI sequence. Participants laid supine in the scanner and could see stimuli and instructions on a back-projection monitor which was placed at a total viewing distance of 108 cm. Before collecting dynamic single-slice data, we started the experiment with the acquisition of static 3D T2 weighted data of the entire head (matrix = 256 × 256 × 250, FOV = 250 × 250 × 250 mm, voxel size = 0.98 × 0.98 × 1.00 mm, TE = 225 ms, TR = 2500 ms, slice thickness = 2 mm, flip angle = 90°, scan duration = 232.5 s), during which the participants were instructed to fixate a dot in the center of the screen. The 3D data was used as a reference for planning the dynamic single-slice data acquisition and to obtain precise knowledge of the eyeball of each participant, which was used later on in the data analysis. Eye movements were then recorded at a temporal resolution of 55.6 ms using a bSSFP sequence in the axial plane (matrix = 224 × 224, FOV = 200 × 200, voxel size = 0.89 × 0.89 mm, TE = 1.28 ms, TR = 2.56 ms, slice thickness = 3 mm, flip angle = 45°, 1020 dynamic scans, total scan duration = 56.7s). To capture eye movements at the highest possible temporal resolution, we used a k-t BLAST factor of 5.

### Experimental Protocol

Our goal was to collect saccades of various amplitudes, in both directions, with and without blinks. Therefore, participants were instructed to continuously look back and forth between two targets along the horizontal meridian. We collected data in 6 separate runs, with target positions either at +/- 2.5°, +/- 5° or +/- 10° and the instruction to make the gaze shift either with or without blinking. In-between runs, which lasted for 56.7 seconds each, we monitored that the lens was fully visible and adjusted the slice position if necessary. Targets were black dots of 0.8° diameter on a grey background. We used MATLAB (The MathWorks, Natick, MA, USA) with the Psychophysics Toolbox [25] for stimuli presentation.

### Data Analysis

#### Preprocessing

We estimated global head motion in the dynamic bSSFP scans using an efficient sub-pixel image registration by cross-correlation algorithm [26]. The coarse location of the eyeball was then identified using the fast-radial symmetry transform [27] and the MR data was cropped around it for further analysis. For better comparison across individual scans and participants, we rescaled the image intensities such that the mean intensity around eyeball center had a value of one for each image.

#### Eye Tracking

Horizontal gaze and retraction, i.e. the amount of translation along the anterior-posterior axis, were quantified for both eyes in each image using the MREyeTrack algorithm [17] with minor modifications (Kirchner2022c). The algorithm segments sclera, lens and cornea in the dynamic MR data by matching the projection of a 3D eyeball model obtained from the static 3D MR data of each participant. Therefore, gaze and retraction are always in reference to the fixation of a target dot in the centre of the screen. For further analysis, we upsampled all eye motion data to a 2 ms time interval and then smoothed the data using a Savitzky-Golay filter of 2nd order and 100 ms window length [28].

#### Saccade detection

Saccades between the two targets were identified using a velocity threshold. If the gaze trajectory passed the midline at a velocity of at least 30°/s this was considered to be part of a saccadic eye movement. On- and offset were then defined as the samples where the velocity fell below 10°/s. Some of the saccades elicited while blinking had dynamic overshoots, which are characterized by exceeding the target followed immediately by a transient return motion in the opposite direction. Typically, saccade kinematics are described by their amplitude (gaze difference between on- and offset), duration and peak velocity. In order to analyze the kinematics of saccades with dynamic overshoots we additionally recorded their maximum excursion (the maximum gaze position during the saccade), overshoot (the gaze difference between maximum excursion and saccade offset) and return (when the gaze was halfway between overshoot and final position).

#### Blink detection

Blinks were detected and classified using eye motion data. They appear as Gaussian-like peaks in the retraction data, so that the retraction velocity profile of each blink is characterized by a positive peak for lid closure and a negative peak for lid opening. We applied a velocity threshold of 1 mm/s to determine on- and offset and further required valid blinks to have a duration of at least 100 ms and a maximum retraction greater than 0.3 mm. The kinematics of blinks were further characterized by their amplitude, the maximum amount of retraction during a blink, and their time of return, which was determined when the retraction was halfway between amplitude and offset.

#### Data exclusion criteria

Precise eye motion estimation relies on high-quality MR images and full visibility of lens and cornea. Even though we corrected the slice position for occasional head motion between runs, out-of-plane head motion within a run could move the lens out of the image. The anterior segment of the eye is also especially sensitive to susceptibility artifacts, which stem from local magnetic field distortions at the interface of air and tissue. If lens or cornea are not well visible, either due to susceptibility artifacts or out-of-plane head motion, this manifests itself as a lower energy functional in the MREyeTrack algorithm. Therefore, we introduced a threshold of 1.5 to the mean energy functionals of sclera, lens and cornea across all runs for each participant (Supplemental Table 1 - https://doi.org/10.6084/m9.figshare.20589888.v1), which excluded the data of participants P1, P3 and P4 from the analysis.

#### Data availability

All data are available from the corresponding author upon reasonable request.

## Acknowledgments

This work has received funding from the European Union’s Horizon 2020 research and innovation programme under the Marie Skodowska-Curie grant agreement No. 734227.

## References

1. Sparks, D. L. The Brainstem Control of Saccadic Eye Movements. Nature Reviews Neuroscience 3, 952–964. doi:10.1038/nrn986 (2002).

2. Demer, J. L. The Orbital Pulley System: A Revolution in Concepts of Orbital Anatomy. Annals of the New York Academy of Sciences 956, 17–32. doi:10.1111/j.1749-6632.2002.tb02805.x (2002).

3. Miller, J. M. Understanding and Misunderstanding Extraocular Muscle Pulleys. Journal of Vision 7, 10.1–15. doi:10.1167/7.11.10 (2007).

4. Quaia, C. & Optican, L. M. Commutative Saccadic Generator Is Sufficient to Control a 3-D Ocular Plant With Pulleys. Journal of Neurophysiology 79, 3197–3215. doi:10.1152/jn.1998.79.6.3197 (1998).

5. Hepp, K. Oculomotor Control: Listing’s Law and All That. Current Opinion in Neurobiology 4, 862–868. doi:10.1016/0959-4388(94)90135-X (1994).

6. Kirchner, J., Watson, T., Bauer, J. & Lappe, M. High-Speed MRI Recordings of Eyeball Lifting, Retraction and Compression during Blinks 2022. doi:10.1101/2022.05.11.491482.

7. Khazali, M. F., Pomper, J. K. & Thier, P. Blink Associated Resetting Eye Movements (BARMs) Are Functionally Complementary to Microsaccades in Correcting for Fixation Errors. Scientific Reports 7, 16823. doi:10.1038/s41598-017-17229-w (2017).

8. Maus, G. W. et al. Target Displacements during Eye Blinks Trigger Automatic Recalibration of Gaze Direction. Current Biology 27, 445–450. doi:10.1016/j.cub.2016.12.029 (2017).

9. Khazali, M. F., Pomper, J. K., Smilgin, A., Bunjes, F. & Thier, P. A New Motor Synergy That Serves the Needs of Oculomotor and Eye Lid Systems While Keeping the Downtime of Vision Minimal. eLife 5, e16290. doi:10.7554/eLife.16290 (2016).

10. Rottach, K. G., Das, V. E., Wohlgemuth, W., Zivotofsky, A. Z. & Leigh, R. J. Properties of Horizontal Saccades Accompanied by Blinks. Journal of Neurophysiology 79, 2895–2902. doi:10.1152/jn.1998.79.6.2895 (1998).

11. Goossens, H. H. & Van Opstal, A. J. Blink-Perturbed Saccades in Monkey. I. Behavioral Analysis. Journal of Neurophysiology 83, 3411–3429. doi:10.1152/jn.2000.83.6.3411 (2000).

12. Goossens, H. H. & Van Opstal, A. J. Blink-Perturbed Saccades in Monkey. II. Superior Colliculus Activity. Journal of Neurophysiology 83, 3430–3452. doi:10.1152/jn.2000.83.6.3430 (2000).

13. Goossens, H. H. & Van Opstal, A. J. Dynamic Ensemble Coding of Saccades in the Monkey Superior Colliculus. Journal of Neurophysiology 95, 2326–2341. doi:10.1152/jn.00889.2005 (2006).

14. Jagadisan, U. K. & Gandhi, N. J. Removal of Inhibition Uncovers Latent Movement Potential during Preparation. eLife 6, e29648. doi:10.7554/eLife.29648 (2017).

15. Gandhi, N. J. & Bonadonna, D. K. Temporal Interactions of Air-Puff-Evoked Blinks and Saccadic Eye Movements: Insights into Motor Preparation. Journal of Neurophysiology 93, 1718–1729. doi:10.1152/jn.00854.2004 (2005).

16. Kirchner, J., Watson, T., Busch, N. A. & Lappe, M. Timing and Kinematics of Horizontal Within-Blink Saccades Measured by EOG. Journal of Neurophysiology 127, 1655–1668. doi:10.1152/jn.00076.2022 (2022).

17. Kirchner, J., Watson, T. & Lappe, M. Real-Time MRI Reveals Unique Insight into the Full Kinematics of Eye Movements. eNeuro 9. doi:10.1523/ENEURO.0357-21.2021 (2022).

18. Enright, J. T. The Aftermath of Horizontal Saccades: Saccadic Retraction and Cyclotorsion. Vision Research 26, 1807–1814. doi:10.1016/0042-6989(86)90132-x (1986).

19. Moon, Y. et al. Positional Change of the Eyeball During Eye Movements: Evidence of Translatory Movement. Frontiers in Neurology 11, 556441. doi:10.3389/fneur.2020.556441 (2020).

20. van Opstal, A. J., Hepp, K., Hess, B. J., Straumann, D. & Henn, V. Two-Rather than Three-Dimensional Representation of Saccades in Monkey Superior Colliculus. Science 252, 1313–1315. doi:10.1126/science.1925545 (1991).

21. Ghasia, F. F. & Angelaki, D. E. Do Motoneurons Encode the Noncommutativity of Ocular Rotations? Neuron 47, 281–293. doi:10.1016/j.neuron.2005.05.031 (2005).

22. Klier, E. M., Meng, H. & Angelaki, D. E. Three-Dimensional Kinematics at the Level of the Oculomotor Plant. Journal of Neuroscience 26, 2732–2737. doi:10.1523/JNEUROSCI.3610-05.2006 (2006).

23. Ramat, S., Leigh, R. J., Zee, D. S. & Optican, L. M. Ocular Oscillations Generated by Coupling of Brainstem Excitatory and Inhibitory Saccadic Burst Neurons. Experimental Brain Research 160, 89–106. doi:10.1007/s00221-004-1989-8 (2005).

24. Schultz, K. P., Williams, C. R. & Busettini, C. Macaque Pontine Omnipause Neurons Play No Direct Role in the Generation of Eye Blinks. Journal of Neurophysiology 103, 2255–2274. doi:10.1152/jn.01150.2009 (2010).

25. Brainard, D. H. The Psychophysics Toolbox. Spatial Vision 10, 433–436. doi:10.1163/156856897X00357 (1997).

26. Guizar-Sicairos, M., Thurman, S. T. & Fienup, J. R. Efficient Subpixel Image Registration Algorithms. Optics Letters 33, 156–158. doi:10.1364/ol.33.000156 (2008).

27. Loy, G. & Zelinsky, A. Fast Radial Symmetry for Detecting Points of Interest. IEEE Transactions on Pattern Analysis and Machine Intelligence 25, 959–973. doi:10.1109/TPAMI.2003.1217601 (2003).

28. Savitzky, A. & Golay, M. J. Smoothing and Differentiation of Data by Simplified Least Squares Procedures. Analytical Chemistry 36, 1627–1639. doi:10.1021/ac60214a047 (1964).

