## Supplemental Table 1 for "Eyeball translations affect saccadic eye movements beyond brainstem control"

| | P1 | P2 | P3 | P4 | P5 | P6 | P7 | P8 | P9 | P10 | P11 |
|---|---|---|---|---|---|---|---|---|---|---|---|
| $E_{sclera}$ | 2.23 | 2.17 | 2.22 | 2.37 | 2.27 | 2.47 | 2.37 | 2.49 | 2.53 | 2.39 | 2.41 |
| $E_{cornea}$ | 0.90 | 2.60 | 1.95 | 0.86 | 2.72 | 2.62 | 2.83 | 1.67 | 1.66 | 2.09 | 2.11 |
| $E_{lens}$ | 2.09 | 2.66 | 1.42 | 2.99 | 2.52 | 2.98 | 2.73 | 2.47 | 2.93 | 2.65 | 2.65 |

**Supplemental Table 1** Data quality assessment for each participant. Individual energy functional of sclera, cornea and lens as determined by normal gradient matching averaged over all bSSFP scans.
